# Unearthing Shifts in Microbial Communities Across a Soil Disturbance Gradient

**DOI:** 10.1101/2021.07.28.454095

**Authors:** Taylor J. Seitz, Ursel M. E. Schütte, Devin M. Drown

## Abstract

Permafrost, an important source of soil disturbance, is particularly vulnerable to climate change in Alaska where 85% of the land is underlain with discontinuous permafrost. Boreal forests, home to plants integral to subsistence diets of many Alaska Native communities, are not immune to the effects of climate change. Soil disturbance events such as permafrost thaw, wildfires, and land use change can influence abiotic conditions which can then affect active layer soil microbial communities. Previously, we found negative effects on boreal plants inoculated with microbes impacted by soil disturbance compared to plants inoculated with microbes from undisturbed soils. Here, we identify the key shifts in microbial inoculant communities altered by soil disturbance using 16S rRNA amplicon sequencing as well as changes in potential functional mechanisms that influence plant growth using long read metagenomics. Across our soil disturbance gradient, microbial communities differ significantly based on the level of soil disturbance. Consistent with previous results, the family *Acidobacteriaceae*, which consists of known plant promoters, was abundant in undisturbed soil, but practically absent in most disturbed soil. In contrast, *Comamonadaceae*, a family with known agricultural pathogens, was overrepresented in most disturbed soil communities compared to undisturbed. Within our metagenomic data, we found that soil disturbance level drives differences in microbial community function. These results indicate that a decrease in plant growth can be linked to changes in the community and functional composition driven by soil disturbance and climate change. Together these results build a genomic understanding of how shifting soil microbiomes may affect plant productivity and ecosystem health as the Arctic warms.

## 1 Introduction

Across the sub-arctic and arctic regions, the warming climate is rapidly affecting Alaska’s ecosystems through shifts in disturbance regimes, including increased fire and permafrost thaw (Chapin et al., 2008; Chasmer and Hopkinson, 2017; Johnstone et al., 2010; Schuur et al., 2015). Boreal forests, which represent 90% of the world’s forests, are a complex mosaic of coniferous trees and peatlands (Wolken et al., 2011). Much of the boreal forests across Alaska are underlain by discontinuous permafrost that is not immune to the pressures of climate change. Permafrost thaw is associated with direct and indirect changes in plant communities due to significant shifts in soil hydrology which in turn affect nutrient availability and carbon dynamics (Inglese et al., 2017; Schütte et al., 2019; Schuur et al., 2007; Sewell et al., 2020; Yang et al., 2013), yet the biotic mechanisms contributing to plant community shifts remain largely unexplored.

In 2011, Mackelprang et al. showed that permafrost thaw leads to rapid changes in community composition and function, including shifts in several genes involving nitrogen and carbon cycling in response to permafrost thaw. They found that during a thaw event, community function within permafrost quickly converged to active layer function (Mackelprang et al., 2011). Consistent with the pattern of permafrost thaw affecting soil microbial communities, in 2017 Inglese et al. found that active layer detachments, a form of permafrost disturbance, significantly affect fungal and Archaeal community composition of the active layer. They observed greater proportions of Nitrososphaerales, an ammonia oxidizing Archaea, in disturbed communities compared to undisturbed, again indicating a shift in nitrogen cycling following permafrost thaw. Within fungal communities, Ingleses et al. (2017) saw a large decrease in Ascomycota in disturbed soils compared to a greater abundance of ericoid mycorrhizal species in undisturbed communities which are often found associated with arctic plant species. They suggest that both the increase of ammonia-oxidizing Archaea and the reduction in hyphal fungi could exacerbate further permafrost thaw and landscape change in the arctic.

While we are gaining more indirect evidence that soil microbial communities are affected by permafrost thaw, it is important to understand changes in the active layer above permafrost where plant-interactions are occurring in the rhizosphere. There is significant evidence that suggests soil microbial communities play a critical role in driving plant community shifts through facilitating nutrient exchange (Bakker et al., 2013), signaling defense or symbiotic responses (Jones et al., 2016), and pathogenic infestations (Hacquard et al., 2017). Increasing our understanding of the relationship between active layer soil microbial communities and soil disturbance events, specifically those resulting in deep permafrost thaw including high temperature wildfires, is crucial in the face of the pressures of climate change in the Arctic. In Seitz et al. (2021), we observed differences in plant growth associated with microbes from a disturbance gradient. Evidence there suggested that microbes from disturbed soils associated with deep permafrost thaw negatively impacted the productivity of several plants commonly found in boreal ecosystems. We found broad scale correlations between the bacterial family *Comamonadaceae* and a decrease in plant growth of plants commonly found in boreal forests.

However, the previous analysis of the soil microbial communities was based on metagenomic sequencing (Seitz et al., 2021), and we wanted to use a measure allowing a closer comparison to other environmental soil microbiome studies (i.e., 16S rRNA amplicon sequencing). Given that studies have demonstrated soil microbial community function shifts with disturbance (Choi et al., 2017; Mackelprang et al., 2011; Santillan et al., 2019), we also aimed to use previously collected metagenomic data to identify any shifts in function that may be leading to the decrease in plant productivity observed in Seitz et al. (2021). To further develop our understanding of the potential impacts of soil disturbance events, including permafrost thaw, we analyzed whether within disturbance level variation can predict the growth of plants from an earlier growth experiment described in Seitz et al. (2021). We hypothesized that within bacterial community diversity will increase with disturbance, and that between community diversity will shift as well. In this study, we evaluated how microbial communities from soils that were subjected to a gradient of physical disturbances were affected based on the intensity of the original disturbance event, and whether microbial community variation can be used to predict plant growth. To do this we generated community composition data which we then tested for shifts in community diversity and composition.

## 2 Methods

### 2.1 Sample Site Description

The soil samples we collected and used for this study were previously described in Seitz et al., 2021. We collected samples from interior Alaska from the Fairbanks Permafrost Experiment Station (FPES; 64.877°N, 147.670°W). This forested site was established and disturbed in 1946 as part of the US Army Corps of Engineers Cold Regions Research and Engineering Laboratory, which aimed to simulate and study how disturbance events influence the ecology of the boreal forest. FPES contains three, 3,721 m^2^ Linell plots, described in Linell et al. (1973) of varying degrees of soil disturbance. When FPES was established, the first plot, referenced as undisturbed (UD), was left untouched. The second plot (semi-disturbed, SD) was cleared of all above ground vegetation and trees, while roots and organic soil layers stayed undamaged. The third plot (most disturbed, MD) was stripped entirely of surface level vegetation and organic material until the mineral soil was reached. Throughout the following 25 years, permafrost in the SD plot degraded to 4.7 m below the surface soils, and the permafrost in the MD plot degraded down to 6.7 m below the surface.

FPES is an example of a Taiga boreal forest found in subarctic regions across the world. The untouched UD plot is characterized by a thick covering of black spruce (*Picea Mariana*) with white spruce (*Picea glauca*) scattered throughout. The UD understory is largely dominated by low-bush cranberry (lingonberry, *Vaccinium vitis-idaea*) and Labrador tea (*Rhododendron groendlandicum*), and the ground is continuously covered with *Sphagnum* and feather moss, and lichen. The SD plot is characterized by a highly mixed stand of black spruce, white spruce, Alaskan birch (*Betula neoalaskana*), and willow (*Salix* spp.). SD trees are taller than those in the MD plot which is characterized by young willow, black spruce, and Alaskan birch (Douglas et al., 2008).

### 2.2 Sample collection and DNA sequencing

To assess how soil microbial communities are shifting based on previous soil disturbance, we collected a total of 48 soil samples on 28 May 2018, with 16 each from UD, SD, and MD plots. The collection of these cores and their original use was described in Seitz et al. (2021). In brief, we extracted cores from a selection of quadrats throughout each disturbance plot to sample the within disturbance level heterogeneity. Using sterile technique, we removed the top layer of moss at each sampling point and then collected the top layer of soil using a 4.5 cm diameter by 10 cm height soil corer. We stored samples in a cooler throughout the duration of sample collection before they were transported back to the lab. After the samples arrived at the lab, we mechanically homogenized each individual soil core and stored them at 4°C.

After homogenizing each soil sample, we extracted DNA from 0.25 g of soil using the Qiagen PowerSoil Kit (Qiagen Inc., USA) according to the manufacturer’s instructions. We then checked the quality and concentration of DNA using a NanoDrop One spectrophotometer (ThermoFisher Science, USA) and a Qubit 4.0 fluorometer (Fisher Science, USA).

### 2.3 PCR and 16S rRNA amplicon sequencing and analyses

After extracting DNA, we performed a 1:10 dilution of all samples to be used for PCR. We amplified the V4 region using dual-indexed 515F (Parada et al., 2016) and 806R (Appril et al., 2015) primers following the EMP PCR protocol (https://earthmicrobiome.org/protocols-and-standards/16s/) to be used for 16S rRNA gene amplicon analysis. Following the first round of amplification, we used gel electrophoresis to determine the presence of PCR products. We ran duplicate PCR reactions and then pooled the two reactions for each unique sample. The amplicons were then sequenced on an Illumina MiSeq using v3 reagents at the Institute of Arctic Biology Genomics Core Lab.

Upon obtaining the 16S rRNA amplicon data, we first demultiplexed all reads using Mr_Demuxy (version 1.2.0, https://pypi.org/project/Mr_Demuxy/1.2.0/). We then processed and analyzed the paired-end, demultiplexed reads using the Quantitative Insights into Microbial Ecology (QIIME2) framework (version 2020.8.0). We used the QIIME 2 plugin DADA2 (https://benjjneb.github.io/dada2/index.html) to obtain a table of representative sequences and frequencies, remove low quality regions of reads, and to merge our paired reads for further processing. We next clustered sequences at 99% identity using the closed reference strategy against the SILVA SSU 138 reference database (Quast et al., 2012) through the VSEARCH plugin. After clustering, we assigned taxonomy with the QIIME 2 taxonomy classifier plugin (Bokulich et al., 2018) using a Naïve Bayes classifier pre-trained on the Silva 138 99% OTUS (515F/806R region). We constructed a rooted phylogenetic tree from representative sequences using the QIIME 2 alignment plug in and then calculated microbial diversity metrics. We next filtered out any chimeric, mitochondrial, and chloroplast reads from our samples using the QIIME 2 plugin feature table.

### 2.4 Amplicon Statistical analyses

All statistical analyses performed in R utilized version 3.6.1. After obtaining our filtered feature table, we used *qiime2R* (version 0.99.35) to import our QIIME 2 data and artifacts into R Studio. We converted our taxonomic data into a *phyloseq* (version 1.30.0) object and filtered out one sample with low sampling depth (less than 14377 reads). We transformed our data into relative abundance measures. We used *vegan* (version 2.5.7, https://github.com/vegandevs/vegan/) to calculate alpha diversity metrics (Pielou’s evenness and Faith’s phylogenetic diversity). We visualized our results with *ggplot2* (version 3.3.3).

To test the significance of overall differences between UD, SD, and MD soil communities, we calculated the Bray-Curtis similarity index to compare community beta diversity using amplicon sequence variants (ASVs). We tested the effects of disturbance on community composition using the Permutational Multivariate Analysis of Variance (PERMANOVA) using the “adonis” function in *vegan* (10,000 permutations). We visualized beta diversity differences using nonmetric dimensional scaling (NMDS) and *ggplot2*.

To visualize community membership across soil samples and taxonomic levels, we generated heatmaps using the R package *gplots* (version 3.1.1). At the phylum level, we filtered out any phyla that were not present in samples at higher than 1% relative abundance. When visualizing at the family level, we filtered out any family that was present at a maximum abundance of less than 5% across all samples.

We generated community functional predictions from the 16S rRNA amplicon data by using Phylogenetic Investigation of Communities by Unobserved States (PICRUSt2) (Langille et al., 2013). We utilized the q2-picrust2 QIIME 2 plugin. We exported the table of KEGG Orthology predictions and imported them into R. We calculated Bray-Curtis distance and ordinated using NMDS in *vegan* to compare predicted community function across disturbance levels.

To examine the relationship between microbial inoculants and resulting plant productivity Seitz et al. (2021) observed in their growth experiment, we utilized linear regression with the *lm* function in R (R Core Team, 2019). Since the first NMDS axis from our 16S rRNA amplicon analysis describes microbial community variation across soil disturbance level, we extracted the scores from the axis of variation, NMDS1 (Figure 2.2), to test whether community variation within disturbance level explains leaf count and plant height from Seitz et al. (2021). For example, for bog blueberry plants, we performed linear regression using MD community variation NMDS scores and plant height as the growth measure. We repeated the regression using SD and then UD scores. We conducted this series of analyses for each plant type and growth measure (height and leaf count). We then adjusted *p*-values using the Benjamini-Hochberg method with a false discovery rate of 0.25 on the basis that follow up studies are relatively low cost. (Benjamini and Hochberg, 1995). We then used *ggplot2* (version 3.3.3) to visualize significant relationships between plant productivity and microbial community variation within each soil disturbance level.

### 2.5 Functional analysis

We performed community functional analyses on previously collected metagenomic data from our soil cores described in Seitz et al. 2021 (ENA Project: PRJEB42020). We utilized the MEGAN-LR (long read) pipeline (Huson et al., 2018) following the parameters described by Arumugam et al. (2019). MEGAN-LR is a tool that performs both taxonomic and functional community classification. Briefly, this pipeline aligns long reads against an NCBI-nr database using DIAMOND (Buchfink et al., 2015) by performing a frame-shift aware DNA-to-protein alignment. The alignments are then processed by MEGAN using an LCA-based algorithm for taxonomic binning and an interval-tree based algorithm for functional binning.

Following functional classification via MEGAN-LR, we utilized the MEGAN comparison function to normalize read counts and compare functions across samples. We then extracted the EGGNOG-COG functions into a matrix. We calculated community diversity metrics using the *vegan* R package (version 2.5.7, https://github.com/vegandevs/vegan/) and visualized beta diversity of COG functions using nonmetric dimensional scaling (NMDS) using *ggplot* (version 3.3.3). We tested the effects of disturbance level through a PERMANOVA using the “adonis” function of *vegan*. To identify potential functional indicators of each disturbance level (UD, SD, MD) we used the R package *indicspecies* (version 1.7.9). In this application, indicator species analysis is using indices of a gene’s or taxon’s (in this case gene) relative abundance and their occurrence to estimate the strength of its association with specific groups (in this instance soil disturbance level). For the indicator analysis, we used only the identified Clusters of Orthologous Groups (COGs) that were identified via MEGAN-LR. Within *indicspecies* we used the function “multipatt” to determine a list of COGs that are significantly associated with each disturbance level. We specified 999 random permutations. We visualized the results using the R package *gplots* (version 3.1.1).

## 3 Results

### 3.1 Microbial community composition differs based on soil disturbance level

We identified a total of 17,005 ASVs within the 47 retained samples with a feature count ranging from 14,076 to 66,994, and a mean feature count of 50,943. After filtering out ASVs that were not seen more than three times in at least 20% of samples, we detected 1005 ASVs. Using the filtered ASV data, we compared both alpha and beta diversity to test for differences in bacterial community across soil disturbance level. We found alpha diversity increased significantly within communities as the level of soil disturbance increased (Figure 1). The comparisons using a Kruskal-Wallis test revealed that the Faith’s Phylogenetic Diversity within MD soil communities was significantly higher than both SD and UD communities (Table 1). We also found that Pielou’s Evenness did not differ significantly between MD and SD communities, but did increase from UD to MD and SD soils.

**Figure 1.**
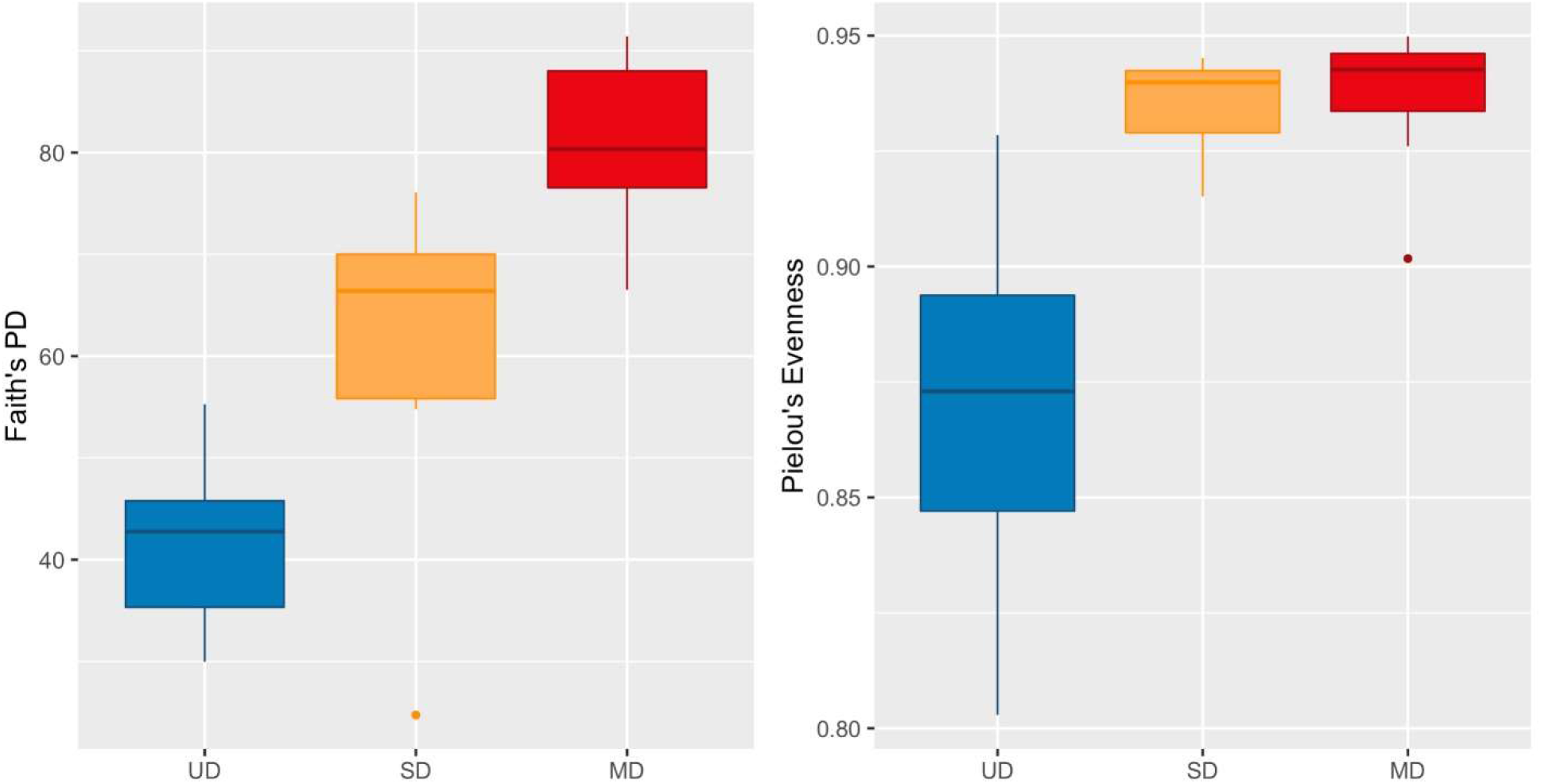
Boxplots of (A) Faith’s Phylogenetic Diversity and (B) Pielou’s Evenness.

We found that microbial community composition differed significantly across the three soil disturbance levels (Figure 2.2; PERMANOVA *F*_2,44_ = 28.682, *p* < .001). Communities cluster based on disturbance level along the first axis. The cluster for MD communities is distinct from both SD and UD communities, while SD and UD communities display some overlap. We did not observe any variation in axis 2 separating the disturbance level groups.

Across all three disturbance levels, we observed shifts in community membership and abundances. We identified 11 phyla that were present at maximum abundances greater than 5% within all samples. Out of those 11 phyla, Proteobacteria was the most abundant phylum, present at mean relative abundances of 40.7% (range 25.2 – 58.8%) in UD, 36.1% (range 27.5 – 43.8%) in SD, and 38.0% (range 29.9 – 46.4%) in MD soil communities. Looking further at the taxonomic composition of the soil communities, we found 17 groups classified to the family level that were present at maximum abundances greater than 5% (Figure 3). Of those, *Acidobacteriaceae* was the most abundant family in UD, present at a mean relative abundance of 16.5% (range 5.46 – 24.7%) compared to a mean of 0.33% (range 0.00 – 3.21%) in MD communities. The most abundant families found in MD communities were *Comamonadaceae* and *Nitrosomonadaceae*, both Proteobacteria, present at mean relative abundances of 3.33% (range 2.0 – 5.7%) and 6.53% (range 3.8 – 10.3%), respectively. Within UD communities *Comamonadaceae* and *Nitrosomonadaceae* were present at mean relative abundances of 0.28% (range 0.03 – 1.57%) and 0.05% (range 0.00 – 0.51%), respectively. We identified uncultured or unclassified taxonomic groupings within all samples. Within the phylum Acidobacteria, we found Subgroup 2 and “uncultured” bacteria, and we also found WD260, a group of uncultured bacteria belonging to the phylum Proteobacteria, to be present at maximum relative abundances greater than 5% throughout our samples.

**Figure 2.**
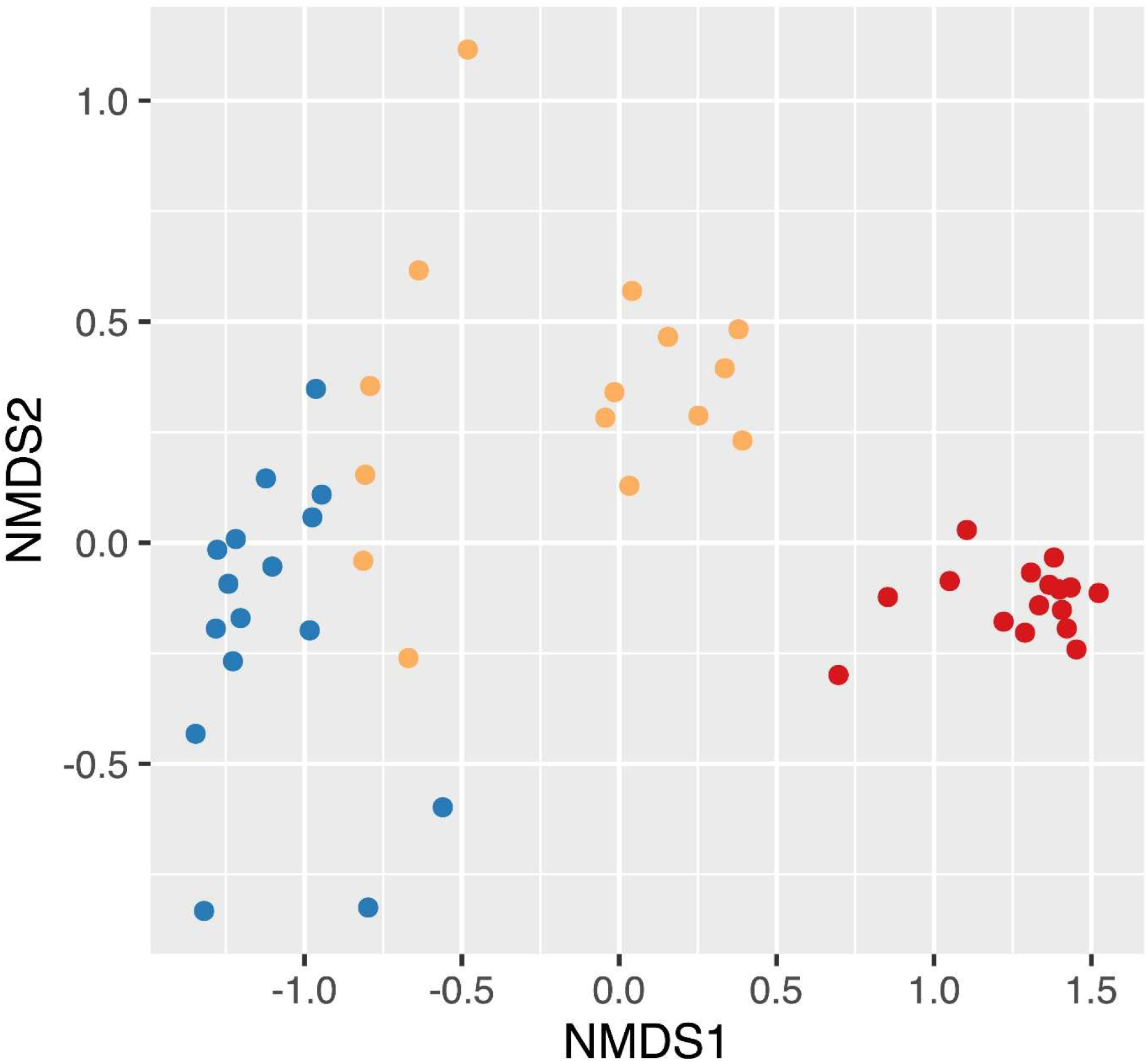
NMDS based on Bray-Curtis dissimilarity distances showing the differences in beta diversity between UD, SD, and MD soil communities. Points colored by the level of FPES soil disturbance with blue = UD (n = 16), gold = SD (n = 15), and red = MD (n = 16).

**Figure 3.**
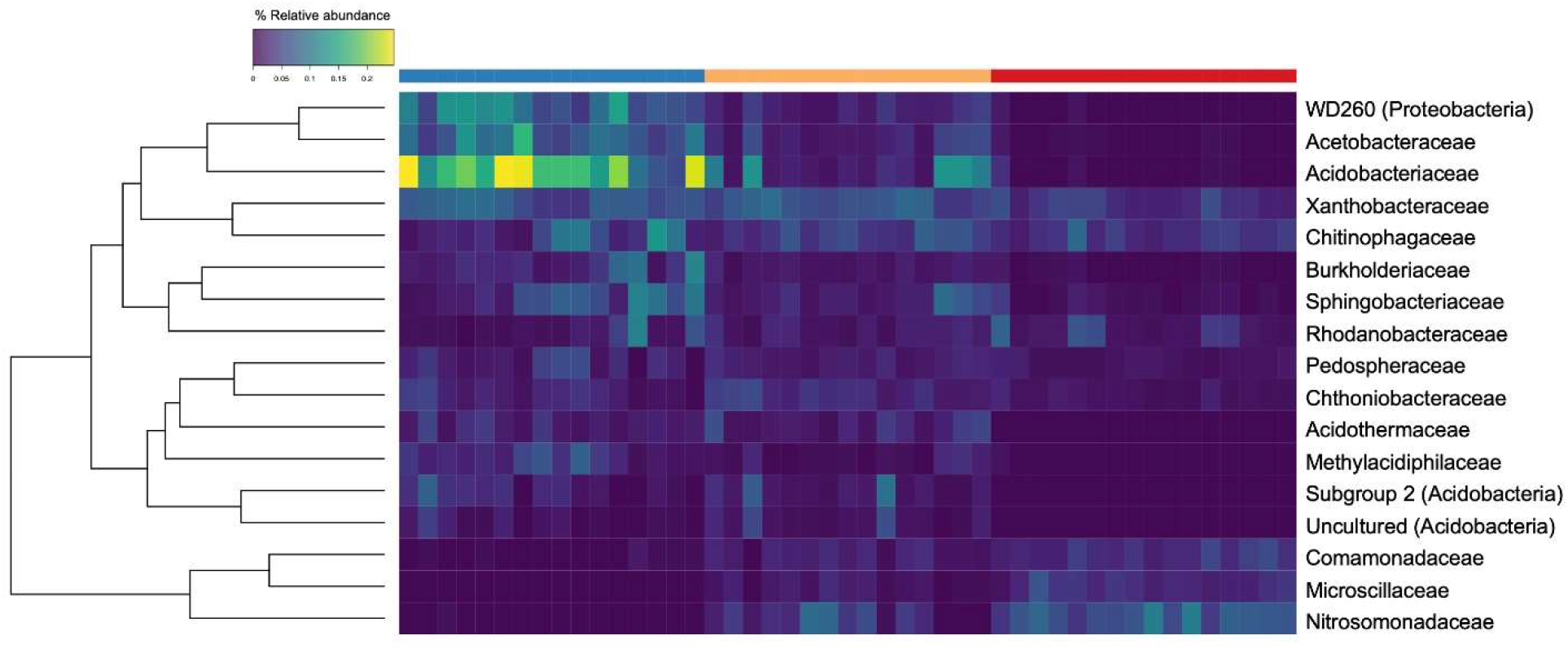
Heatmap of bacterial families present at maximum abundances greater than 5%. Each row corresponds to a family, and each column corresponds to an individual soil core community. The top row signifies the level of soil disturbance of each core with blue = UD (n = 16), gold = SD (n = 15), and red = MD (n = 16). The rows are clustered by Bray-Curtis distance and the color of each box displays the relative abundance of the taxon within that core.

### 3.2 Microbial community variation as a predictor for plant growth

Following observing significant changes in microbial community diversity depending on the level of FPES soil disturbance, we sought to determine whether within disturbance level community variation could predict the growth of plants from an earlier experiment described in Seitz et al. (2021). Within each disturbance level we regressed the productivity measures (height or leaf count) against the NMDS1 values for each corresponding microbial community used as an inoculant in Seitz et al., 2021. We observed significant negative linear relationships between MD microbial community variation and plant growth within bog blueberry height, low-bush cranberry height and leaf count, and Labrador tea height and leaf count (Table 2; Figure 4: A, D, E, and F). Within disturbance level variation was present for bog blueberry, low-bush cranberry, and Labrador tea plants grown with MD microbes, for as microbial communities became more extreme and more positive along the NMDS1 axis (NMDS1 > 0.5), plant growth continued to decrease. Variation within SD and UD communities was not a significant predictor of height or leaf count in any plant species except for black spruce (height) from Seitz et al. (2021) (Table 2.2).

**Figure 4.**
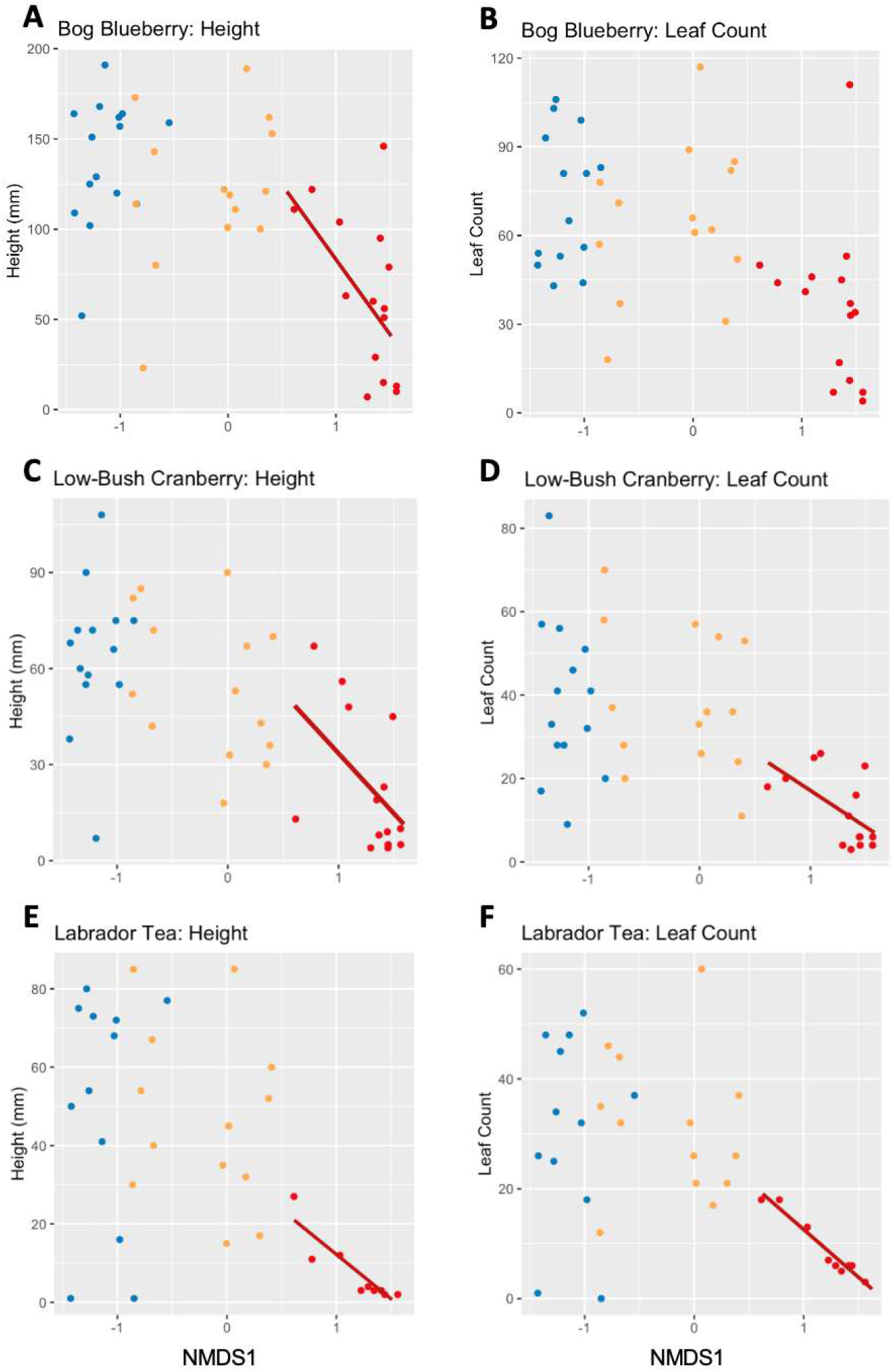
Scatterplots showing the effects of microbial community variation and plant productivity (16S rRNA amplicon NMDS1 and height/leaf count) of bog blueberry (A, B), low-bush cranberry (C, D), and Labrador tea (E, F). Each point represents a soil community, colored by the level of FPES soil disturbance with blue = UD (n = 16), gold = SD (n = 15), and red = MD (n = 16). Solid lines represent a significant linear relationship.

**Table 2.1.**
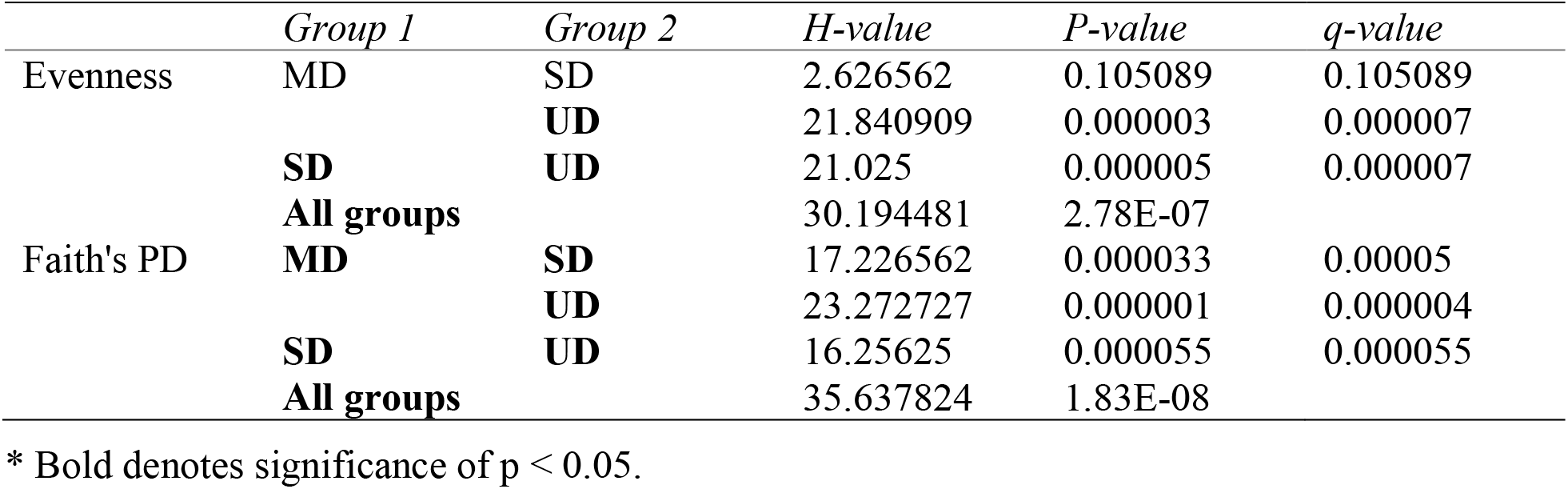
All groups and pairwise comparison of alpha diversity by Kruskal-Wallis.

**Table 2.2.** Results from linear models of plant growth measures predicted by 16S NMDS1.

| **Plant** | **FPES level** | **Growth measure** | **Estimate** | **Standard Error** | **t value** | **p value** | **BH adjusted p value** |
| --- | --- | --- | --- | --- | --- | --- | --- |
| BB | UD | height | 47.4 | 39.0 | 1.21 | .246 | .486 |
|  | SD | height | 27.0 | 23.0 | 1.17 | .264 | .486 |
|  | **MD** | **height** | **-83.5** | **36.7** | **-2.27** | **.041** | **.244** |
|  | UD | leaf count | 13.4 | 36.4 | 0.37 | .719 | .719 |
|  | SD | leaf count | 14.6 | 14.3 | 1.02 | .329 | .486 |
|  | MD | leaf count | -21.9 | 25.5 | -0.86 | .405 | .486 |
| CB | UD | height | 23.8 | 37.9 | 0.63 | .542 | .542 |
|  | SD | height | -17.6 | 12.1 | -1.45 | .172 | .344 |
|  | **MD** | **height** | **-38.5** | **18.6** | **-2.07** | **.06** | **.181** |
|  | UD | leaf count | -26.2 | 30.7 | -0.85 | .411 | .493 |
|  | SD | leaf count | -9.6 | 9.5 | -1.02 | .33 | .493 |
|  | **MD** | **leaf count** | **-17.3** | **7.0** | **-2.48** | **.029** | **.175** |
| LT | UD | height | 4.2 | 36.0 | 0.12 | .909 | .948 |
|  | SD | height | -11.0 | 13.4 | -0.82 | .429 | .858 |
|  | **MD** | **height** | **-23.4** | **4.2** | **-5.57** | **.001** | **.003** |
|  | UD | leaf count | 1.4 | 21.2 | 0.07 | .948 | .948 |
|  | SD | leaf count | -3.5 | 8.0 | -0.44 | .669 | .948 |
|  | **MD** | **leaf count** | **-17.4** | **1.6** | **-11.14** | **< .001** | **< .001** |
| BS | **UD** | **height** | **37.7** | **16.6** | **2.27** | **.04** | **.238** |
|  | SD | height | -7.3 | 8.3 | -0.88 | .396 | .756 |
|  | MD | height | 7.3 | 11.7 | 0.62 | .543 | .756 |
|  | UD | leaf count | 70.8 | 47.3 | 1.50 | .156 | .469 |
|  | SD | leaf count | -10.0 | 27.1 | -0.37 | .721 | .756 |
|  | MD | leaf count | 18.7 | 59.2 | 0.32 | .756 | .756 |
| FW | UD | height | 3.4 | 25.4 | 0.13 | .896 | .896 |
|  | SD | height | -31.6 | 27.6 | -1.14 | .277 | .613 |
|  | MD | height | 9.4 | 22.6 | 0.42 | .682 | .833 |
|  | UD | leaf count | -6.2 | 5.8 | -1.07 | .306 | .613 |
|  | SD | leaf count | -1.7 | 4.2 | -0.40 | .694 | .833 |
|  | MD | leaf count | -6.2 | 3.4 | -1.82 | .091 | .545 |

\*Bold denotes significant Benjamini-Hochberg (BH; Benjamini and Hochberg, 1995) adjusted p value < 0.25.

**Table 2.3.** Highly sensitive and specific indicator functions from EggNOG annotations.

| **FPES** | **COG** | **Sensitivity (A)** | **Specificity (B)** | **Statistic** | **P value** |
| --- | --- | --- | --- | --- | --- |
| UD | COG4845 | 0.8029 | 0.875 | 0.838 | 0.001 |
| SD | COG5163 | 0.9804 | 0.5 | 0.7 | 0.001 |
| SD | COG5033 | 0.852 | 0.5 | 0.653 | 0.001 |
| MD | COG3940 | 0.9239 | 0.625 | 0.76 | 0.001 |
| MD | COG5598 | 0.8921 | 0.75 | 0.818 | 0.001 |
| MD | COG1423 | 0.88 | 0.5625 | 0.704 | 0.001 |
| MD | COG4305 | 0.8765 | 0.5625 | 0.702 | 0.001 |
| MD | COG1908 | 0.8745 | 0.6875 | 0.775 | 0.001 |
| MD | COG4559 | 0.8584 | 0.625 | 0.732 | 0.001 |
| MD | COG1140 | 0.8409 | 0.75 | 0.794 | 0.001 |
| MD | COG3487 | 0.8401 | 0.625 | 0.725 | 0.002 |
| MD | COG1148 | 0.8199 | 0.8125 | 0.816 | 0.001 |
| MD | COG4452 | 0.806 | 0.5 | 0.635 | 0.007 |
| MD | COG2231 | 0.8051 | 0.5 | 0.634 | 0.009 |

### 3.3 Differences in community function based on soil disturbance level

To compare the diversity of community functions across the three levels of soil disturbance at FPES, we first inferred gene function from the 16S rRNA amplicons using PICRUSt2. We found that predicted community function varies significantly based on the FPES disturbance level, with MD communities clustering distinctly from the SD and UD communities along the first axis (Figure S1, PERMANOVA *F*_2,45_ = 0.17433, *p* = .001). To test this prediction, we compared the observed EggNOG (evolutionary genealogy of genes: Non-supervised Orthologous Groups) annotations from our metagenomic data. We identified 2891 clusters of orthologous genes (COGs) and 7,980 non-supervised orthologous groups across all samples. Using Bray-Curtis distances and a PERMANOVA analysis, we found that community function varies significantly based on the level of soil disturbance (Figure 5, PERMANOVA *F*_2,44_ = 0.17431, *p* = .001).

**Figure 5.**
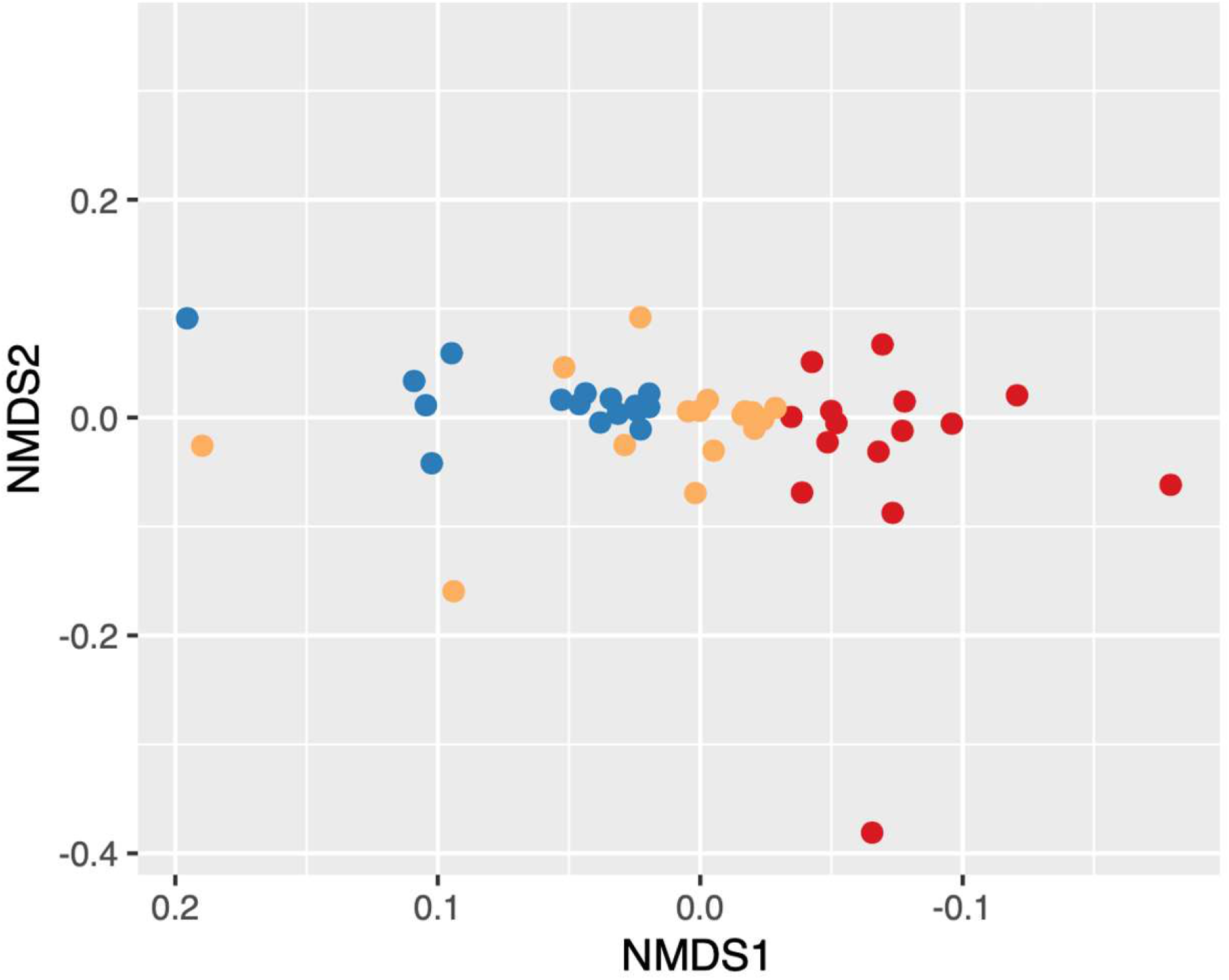
NMDS of Bray-Curtis dissimilarity distances of community function. Functions based on MEGAN-LR EGGNOG COG classifications. Points colored by the level of FPES soil disturbance with blue = UD (n = 16), gold = SD (n = 15), and red = MD (n = 16).

To identify specific functions that are characteristic of the three levels of soil disturbance, we performed an indicator analysis on the observed genomic functions of our microbial communities. The indicator analysis revealed a total of 74 significant COGs across all three FPES levels, with 45 MD, 17 SD, and 12 UD indicators. After further filtering for a higher specificity (A ≥ 0.8) and sensitivity (B ≥ 0.5), we found 14 COG indicators, with 11 belonging to MD, two SD, and one UD (Figure S2, Table 3). Within the indicator COGs we identified a variety of clusters including those encoding for a cell-wall degrading enzyme, disulfide reductases, and a nitrate reductase.

## 4 Discussion

Supporting our hypothesis that soil disturbance drives community shifts, we found that bacterial community composition significantly differed with level of soil disturbance (Figures 1 and 2). We observed low within disturbance level diversity and strong structure of communities between levels. We also detected significant increases in alpha diversity with increases in soil disturbance. This is a phenomenon that is observed within many ecosystems subject to many different types of disturbance, suggesting that disturbance promotes diverse communities due to the creation of new niches within the environment (Galand et al., 2016; Violle et al., 2010).

Our findings of microbial community diversity increasing with disturbance are consistent with long-term (Feng et al., 2020) and short-term (Luláková et al., 2019) studies of the effects of warming soils and permafrost thaw on active layer bacterial communities. Feng et al. (2020) and Luláková et al. (2019) both saw alpha diversity measures significantly increase as topsoil temperatures increased. This trend of alpha diversity increasing along with soil disturbance has also been demonstrated in soils from a variety of land and forest types that have been subject to physical disturbances such as logging, agricultural conversion, and fires (Chatterjee et al., 2008; de Carvalho et al., 2016; Shen et al., 2016; Zhou et al., 2017). The physical soil disturbance and resulting deep permafrost thaw that the MD level plot underwent has created a heterogenous landscape (Douglas et al., 2008), which may be contributing to this increased diversity by affecting the ecological niches and competition ability of bacteria for resources, similar to what can occur after forest fires (Shen et al., 2016).

The lower community evenness that we observed within our UD communities may be due to the high relative abundances of Acidobacteria, compared to our MD and SD communities. When ASVs were grouped at the family level, *Acidobacteriaceae* was the most abundant bacterial family within the UD communities, compared to being nearly absent in MD communities. Members of *Acidobacteriaceae* are typically found to be present at higher abundances in acidic soils that are carbon poor (Kielak et al., 2016; Osburn et al., 2019). In contrast, *Nitrosomonadaceae*, a copiotrophic bacterial family of ammonia oxidizers, was found to be the most abundant family within the MD soil communities, followed by *Comamonadaceae. Nitrosomonadaceae* are ammonia oxidizing bacteria (AOB) that are commonly found in carbon-rich environments such as wastewater and soils (Prosser et al., 2014). An increase of copiotrophic bacteria with disturbance is consistent with the body of literature (Masse et al., 2017; Mickan et al., 2018; Osburn et al.,2019). Masse et al. (2017) previously detected that oligotrophic bacteria, such as *Acidobacteriaceae*, were more abundant in natural boreal forest soils, compared to soils that were reclaimed or reconstructed after disturbance which displayed higher abundances of copiotrophic bacteria.

Connecting back to our previous work described in Seitz et al. (2021), we found that the variation in microbial communities across soil disturbance is a significant predictor of plant productivity measures for bog blueberry (*Vaccinium uliginosum*), low-bush cranberry *(Vaccinium vitis-ideae*), and Labrador Tea (*Rhododenron groenlandicum*) plants. Specifically, we can predict that plants inoculated with a microbial community indicative of the most extreme disturbance, will demonstrate decreased growth compared to plants inoculated with communities from less disturbed soils. This significant relationship between microbial community variation and plant productivity was observed within plants inoculated with MD microbes, whereas we saw no significant relationships between plants inoculated with SD or UD microbes and community variation. This suggests that there is a threshold effect occurring across the disturbance levels, and that soil microbial communities subject to disturbance resulting in permafrost thaw may not have the capacity to influence plant productivity of these boreal species until affected by an event of large enough magnitude. This microbial threshold we are observing could potentially be linked to a disturbance threshold of deep permafrost thaw. Over the past 20 years, researchers have observed accelerated permafrost thaw within the northern boreal Taiga plains likely due to a combination of hydro-climactic changes and increases in annual air temperatures (Chasmer and Hopkinson, 2016; Jorgenson et al., 2006; Lara et al., 2015). As permafrost thaw is accelerated and other disturbance events are increasing in the artic, we may start to observe the effects through changes in soil microbial communities and plant productivity.

Following analysis using the hypothesis generating tool, PICRUSt2, we predicted that as microbial communities shift with disturbance, so will the observed functional community. Using the metagenomic sequencing data, we identified functional indicators across all our FPES sites representative of each level of soil disturbance, with most indicators belonging to MD soils. The gene cluster with the highest specificity within MD is COG3940, which encodes for alpha-N-arabinofurandase (ABF). ABFs are mainly extracellular enzymes that degrade lignocelluloses and cell walls (De loannes et al., 2000; Numan and Bhosle, 2006), and have been shown to play a role in triggering plant immune responses. In 2016, Wu et al. found that a novel ABF plays a critical role in the pathogenicity of the fungal pathogen, *Magnaporthe oryzae*, that causes rice blast disease. The secreted ABF degrades cell walls, further leading to a decrease in productivity in infected plants (Wu et al., 2016). The presence of a gene cluster encoding for alpha-N-arabinofurandase in our MD communities provides a possible mechanism behind the decrease in plant productivity demonstrated in Seitz et al. (2021), yet further studies involving gene expression and enzyme activity would be required to identify if this is part of a mechanism at play.

The critical gene involved in ammonia oxidization (amoA; Bárta et al., 2017; Prosser et al., 2014) was present across all our sample communities, however this gene cluster was not identified as a functional indicator of MD communities where *Nitrosomonadaceae* were very abundant. Curiously, we identified COG1140, a gene cluster encoding for a nitrate reductase beta subunit, to be a highly specific and significant indicator of MD soil communities. Nitrate losses within soils have long been known to follow disturbance events in forests such as fire, disease, or clear cutting (Hobara et al., 2001; Neary et al., 1999; Pardo et al., 2002; Vitousek et al. 1979). This potential loss of available nitrate could be part of the mechanism at play in the reduced plant productivity that Seitz et al. (2021) observed within plants grown in MD community inoculated soils.

As climate change and anthropogenic-driven soil disturbance events including wildfires (Calef et al., 2008; Chapin et al., 2008; Partain et al., 2016) and permafrost thaw (Chasmer and Hopkinson, 2016; Jorgenson et al., 2006; Lara et al., 2015) are increasing across the north, we need to continue to study how disturbance influences the microbiomes of active layer soils that are both directly and indirectly involved in mediating plants within boreal forests. In this study, we show that active layer soil microbial communities are influenced by the initial disturbance event and permafrost thaw at FPES, however we cannot conclude which specific environmental factors are associated with community shifts. We observed significant shifts in microbial community variation, as well as microbial community function. Our evidence for microbial community variation being a significant predictor of plant productivity in response to soil disturbance and permafrost that suggests that this effect will occur once a threshold level of disturbance is reached. These consistent effects of disturbance on active layer microbial communities, combined with the knowledge of how they in turn can affect plant productivity (Seitz et al., 2021), highlight the need for more research into how on-going and future disturbance events will influence the ecological dynamics in boreal forests.

## 5 Acknowledgements

We would like to thank Dr. Anne-Lise Ducluzeau and Tracie Haan for laboratory support, as well as Scout McDougall and Jennie Humphrey for support with data collection. We would also like to thank Drs. Christa Mulder and Brandon Briggs for their feedback and guidance on analysis and writing. Thanks to Dr. Tom Douglas from the Cold Regions Research and Engineering Laboratory (CRREL)-Alaska for providing access to the field site. This research was supported by the Alaska BLaST program, the Institute of Arctic Biology, and Alaska INBRE. Research reported in this publication was supported by an Institutional Development Award (IDeA) from the National Institute of General Medical Sciences of the National Institutes of Health under grant number 2P20GM103395 as well as under three linked awards number RL5GM118990, TL4GM118992 and 1UL1GM118991.

## 9 Supplementary Material

**Figure S1.**
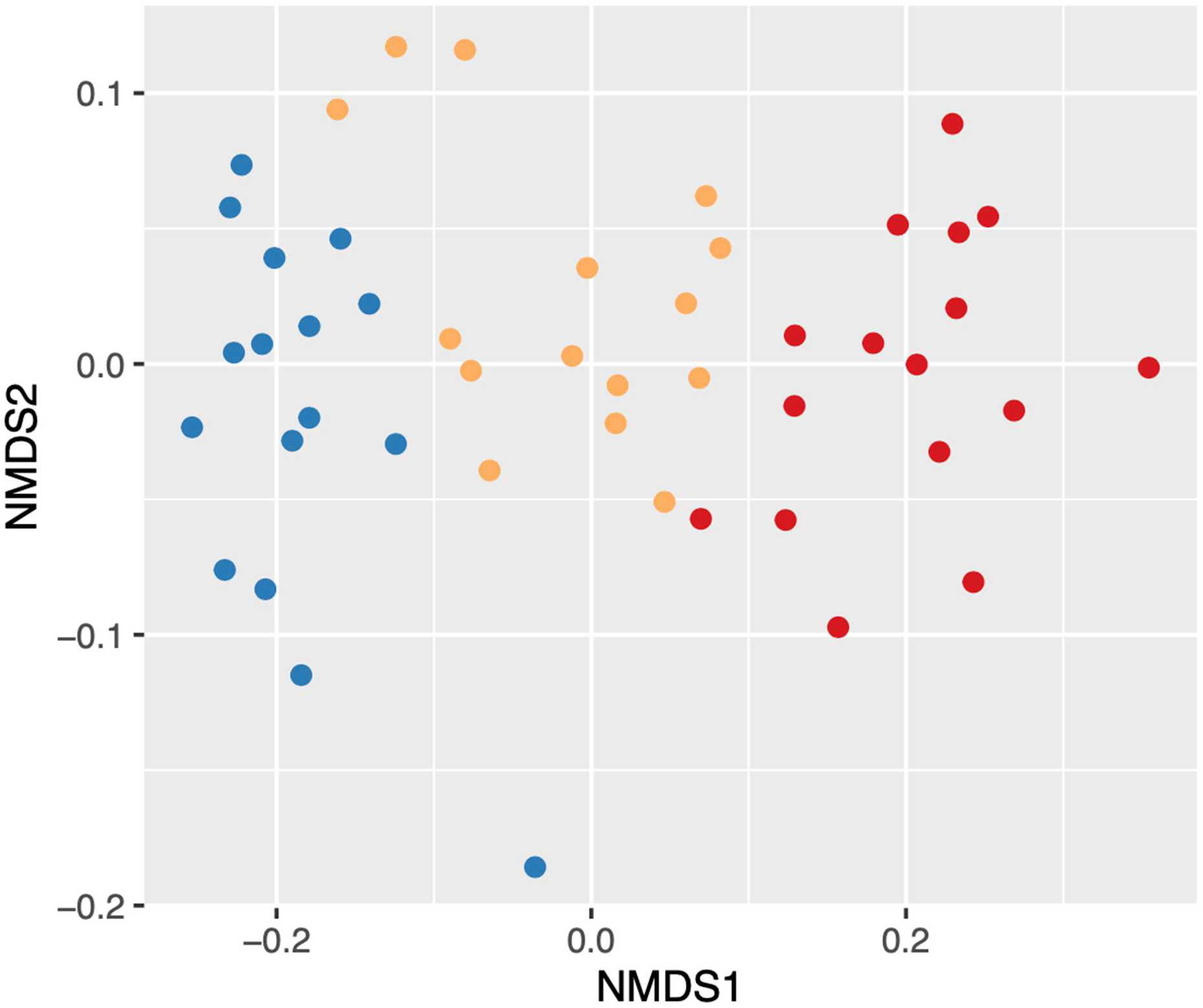
NMDS of Bray-Curtis dissimilarity distances of predicted KEGG KO community function via PICRUSt2 analysis.

**Figure S2.**
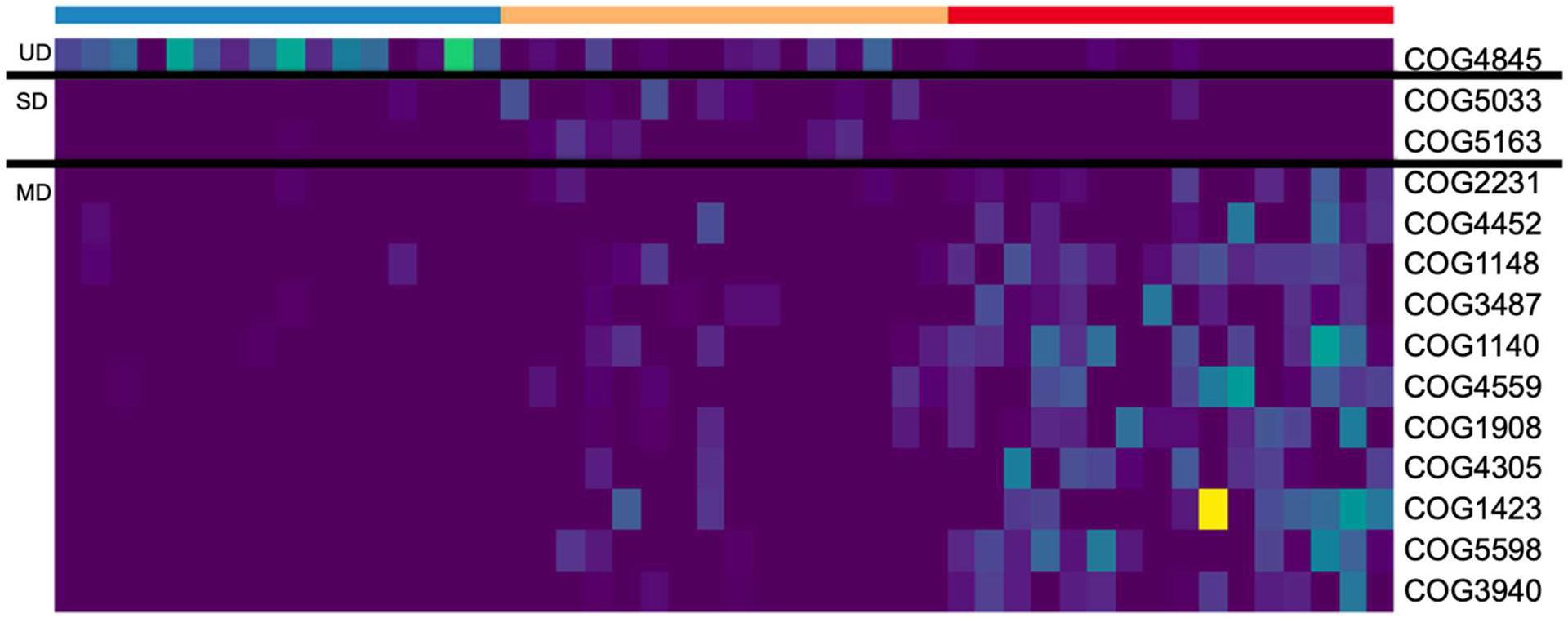
Heatmap of highly sensitive and specific indicator functional EggNOG annotations from metagenomic data. Each row corresponds to a COG, and each column corresponds to an individual soil core community. The top row signifies the level of soil disturbance of each core with blue = UD (n = 16), gold = SD (n = 15), and red = MD (n = 16). The color of each box corresponds to the normalized read count of the COG within each core.

